# ProPicker: Promptable Segmentation for Particle Picking in Cryogenic Electron Tomography

**DOI:** 10.1101/2025.02.27.640512

**Authors:** Simon Wiedemann, Zalan Fabian, Mahdi Soltanolkotabi, Reinhard Heckel

## Abstract

Cryogenic electron tomography (cryo-ET) produces detailed 3D images (tomograms) of cellular environments. A key step in cryo-ET data analysis is detecting all instances of a specific particle across tomograms (particle picking). This is a challenging object detection task due to strong noise, artefacts, and the crowded cellular context. Here, we propose ProPicker, a pretrained, promptable 3D segmentation model that enables a flexible and data-efficient particle picking workflow. By specifying a prompt, ProPicker is conditioned to detect a particle of interest and can then be used directly or can be fine-tuned to a particle-specific picker for improved accuracy. Experiments on simulated and real-world tomograms show that, using a single prompt, ProPicker achieves performance close to or on par with state-of-the-art methods while being up to an order of magnitude faster. Moreover, ProPicker can detect particles not seen during training. Fine-tuning ProPicker outperforms state-of-the-art particle-specific pickers if limited training data is available.

## 1 Introduction

Cryo-electron tomography (cryo-ET) is gaining popularity due to its unique ability to image biological macromolecules in their native environments [TB20; HS21]. An ambitious goal of cryo-ET is to obtain an atlas of the cell with all of its constituent macromolecules mapped in their native environment. This would revolutionize our understanding of essential protein interactions and has the potential to provide breakthroughs in modern medicine spanning cell biology to drug discovery [Bod+23].

In this paper, we focus on particle picking, which is the task of finding all instances of a particle of interest in 3D volumes, called tomograms, obtained with cryo-ET. Particle picking is an essential step and often a bottleneck in important cryo-ET analysis pipelines [Gen+23].

Particle picking is a 3D object detection problem and is particularly challenging for various reasons. Due to the fundamental limitations of data acquisition in cryo-ET, tomograms have a very low signal-to-noise ratio and exhibit strong artefacts. Moreover, tomograms are often large (200 × 1000 × 1000 voxels and larger), and cryo-ET datasets can consist of hundreds of tomograms, making their analysis computationally demanding [Gen+23; Zen+23]. Finally, due to the significant diversity in protein types within the cell, there is a vast array of unique object classes to be detected, many of which only differ subtly, rendering differentiation challenging. For instance, the human body alone is estimated to contain more than 20,000 unique proteins [LB21].

A particle picking method should be fast and flexible, i.e., should be able to accurately pick any particle of interest using no or little additional data for, e.g., model training or fine-tuning. Existing methods for particle picking are either slow or not flexible. Most state-of-the-art methods [Moe+21; DTT+23; Liu+24] are based on deep learning models which can detect only a small, fixed set of particles and require training on large amounts of labeled data, which is particularly difficult to obtain in the cryo-ET domain.

Here, we propose ProPicker, a **Pro**mptable particle **Picker** that can target any particle of interest via a versatile prompting mechanism. ProPicker is trained on a large and diverse synthetic dataset and leverages an efficient 3D segmentation network to segment particles of interest in tomograms and to accurately locate their positions. The promptable design enables the user to select which single particle class is targeted by the segmentation model (prompt-based picking). Particle-specific fine-tuning can be used to improve on the out-of-the-box performance of ProPicker. The promptable design of ProPicker is inspired by methods for the segmentation of arbitrary (natural) 2D images like the Segment Anything Model (SAM) [Kir+23] and CLIPSeg [LE22].

ProPicker is able to pick new particles unseen during training in synthetic tomograms with high F1 scores based on a single prompt (see Section 4.1.1). Experiments on two challenging real-world datasets, `DS-10440` [Pec+24] and `EMPIAR-10988` [DTT+23], show that prompt-based picking can detect large particles and such that produce strong contrast, e.g., ribosomes and apoferritin.

In both synthetic and real-world settings, we also encountered situations where prompt-based picking with ProPicker and other flexible baselines, i.e., TomoTwin [Ric+23] and CryoSAM [Zha+24] do not give satisfactory results. In Section 4.1.2 and Section 4.2, we discuss such scenarios and demonstrate that fine-tuning ProPicker on little data (even *≤* 25% of a tomogram) can improve the F1 score by factors of up to 4, depending on the particle.

Furthermore, we show that fine-tuning ProPicker requires less data than training the state-of-the-art particle-specific method, DeepETPicker [Liu+24], to achieve comparable picking performance.

Finally, ProPicker is the fastest among flexible pickers capable of detecting particles based on a prompt, being up to an order of magnitude faster than the state-of-the-art TomoTwin [Ric+23] (see Section 4.1.1). Our findings illustrate the promising capabilities of ProPicker, and highlight the need for large, diverse,and extensively annotated real-world training datasets to unlock its full potential (see Section 5).

## 2 Background & Related Work

Deep learning methods have already revolutionized particle picking from 2D micrographs in the context of single particle cryo-EM [Wan+16; Bep+19; Wag+19]. They are also on the rise for picking particles in 3D tomograms produced with cryo-ET, but traditional methods still play an important role:

### Particle Picking with Template Matching

Template Matching is the most widely used traditional method for particle picking in cryo-ET [Boh+00; CL+24]. Template-matching-based methods compare a template of the particle to be picked to candidate sub-tomograms extracted by a 3D sliding window. This approach is flexible, as it can target any particle as long as a template is available. However, template matching is computationally demanding (up to several hours per tomogram), as the stride of the sliding window needs to be small for accurate picking [Gen+23; MSK24].

Building upon classical template-based approaches, the recent TomoTwin method [Ric+23] utilizes a learned convolutional encoder to map both template and sub-tomogram into a structured latent space, where similarity is evaluated. The training of the encoder follows a deep metric learning approach such that latent representations of particles of the same class have high cosine similarity, while those of different classes have low cosine similarity. TomoTwin can outperform classical template matching in terms of speed and performance, and is more convenient to use [Ric+23].

### Particle Picking with Deep Learning-Based Object Detection

Particle pickers using deep learning-based object detection often outperform template matching in terms of performance and picking speed [Gub+20; Gen+23]. Many such pickers use a convolutional network to segment particles of interest belonging to one or more classes and produce candidate particle locations by clustering the predicted segmentation masks. Examples for such methods include DeepFinder [Moe+21], DeePiCt [DTT+23], and DeepETPicker [Liu+24]. Deep learning-based object detection approaches for particle picking are typically significantly faster than template-matching-like methods [Gub+20]. However, they are trained on datasets containing a fixed set of particles of interest and are therefore limited to picking particles seen during training.

As a first step towards an entirely training-free segmentation-based picker, Zhao et al. [Zha+24] recently proposed CryoSAM. CryoSAM takes center coordinates of target proteins as input. Leveraging DINOv2 [Oqu+23], it extracts features around these points and generates new prompts via feature matching with other parts of the tomogram. Based on newly generated prompts, the 2D segmentation model SAM [Kir+23] is then used to segment the particles.

### Relation of ProPicker to Existing Particle Pickers

ProPicker shares the setup of picking particles based on a single observation (without any training or fine-tuning) with TomoTwin and CryoSAM. In contrast to TomoTwin, ProPicker does not rely on slow template matching, but is based on fast 3D segmentation. CryoSAM is segmentation-based, but the technique used differs significantly from ProPicker: CryoSAM does not involve any training on cryo-ET specific data, but relies on 2D models trained in the general domain on mostly natural images. ProPicker on the other hand is a 3D segmentation model trained on domain-specific data.

The second application of ProPicker, i.e., fine-tuning ProPicker on annotated data to pick a specific particle, assumes the same problem setup as deep-learning-based pickers which involve training a particle-specific segmentation network, e.g., DeepETPicker [Liu+24].

## 3 ProPicker: Promptable Segmentation for Particle Picking

After formally stating the particle picking problem, we describe ProPicker’s promptable segmentation model, and detail how to use the trained model for prompt-based particle picking and fine-tuning.

### Problem statement

ProPicker takes a tomogram ***x*** *∈* ℝ_*n*×*n*×*n*_, and a prompt ***p*** *∈* ℝ_*m*×*m*×*m*_ representing the particle of interest as input. The output is a finite set **C* ⊂ {*1, …, *n}*_3_ that contains predicted coordinates (voxel indices) of particle centers.

### Promptable segmentation model

The ProPicker model shown in Figure 1 consists of a prompt encoder and a conditional segmentation model. The prompt encoder *ε* : ℝ_*m*×*m*×*m*_ *→*ℝ_*d*_ extracts a feature vector *ε* (***p***) *∈* ℝ_*d*_ that encodes information required for efficiently detecting the particle in tomograms.

**Figure 1:**
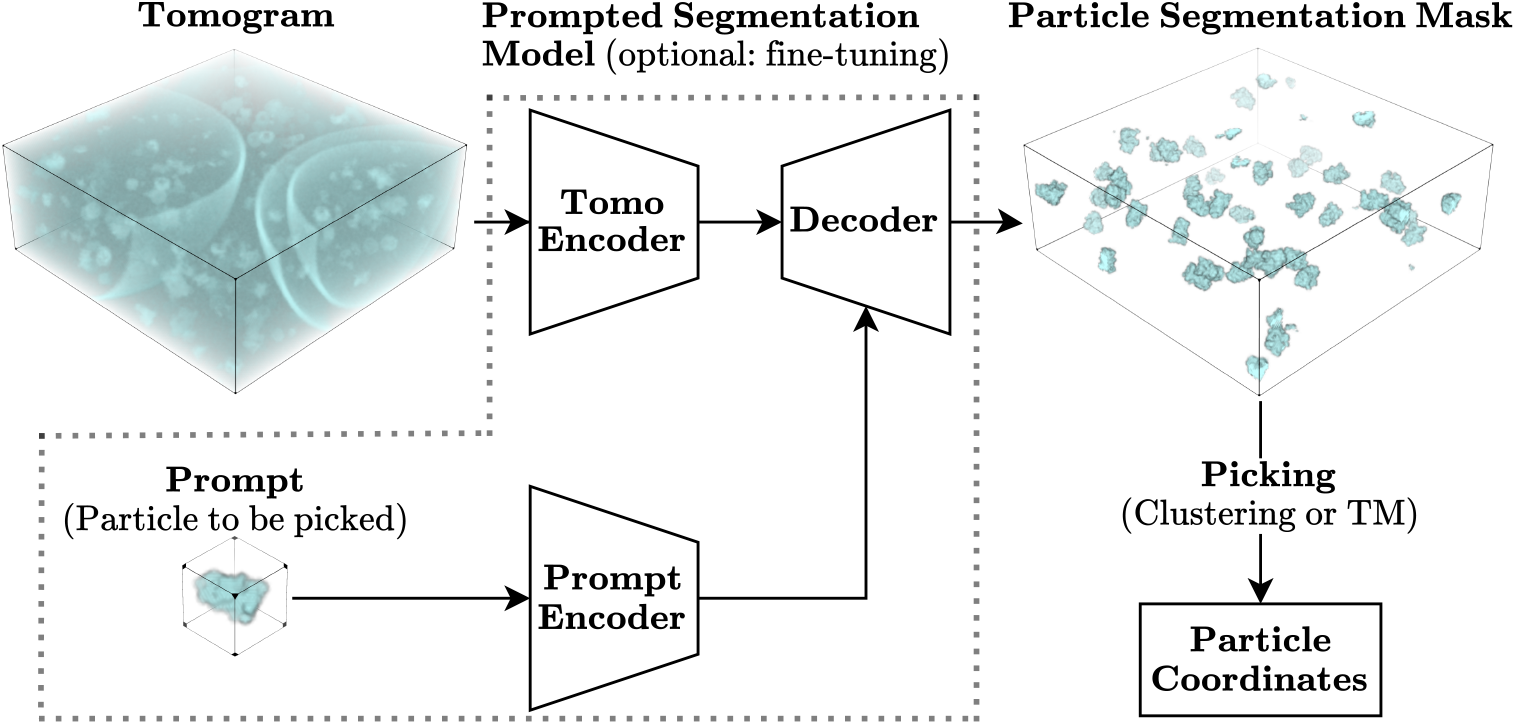
Overview of ProPicker: First, ProPicker extracts features of the particle to be picked in the tomogram with a prompt encoder. Conditioned on the features, ProPicker segments the tomogram with a pre-trained segmentation model. Finally, the the particle coordinates are extracted from the output segmentation mask. The conditioned segmentation model can be fine-tuned on annotated data. The 3D objects in this figure have been rendered using data from the SHREC 2021 dataset [Gub+20].

Given an input volume ***x*** *∈* ℝ^*n*×*n*×*n*^ and prompt ***p***, our promptable segmentation model *S* : ℝ^*n*×*n*×*n*^ ×ℝ^*d*^ *→* R^*n*×*n*×*n*^ can be conditioned on the input prompt in order to steer the output map ***y*** *∈* [0, 1]^*n*×*n*×*n*^ to the desired particle class, that is ***y*** = **S*(* ***x***; *ε* (***p***)).The model output ***y*** is the voxel-wise prediction of the model with respect to the absence/presence of the particle described in the input prompt. We detail the architecture of the conditional segmentation model in Appendix A.

We train the promptable segmentation model on a synthetic dataset of tomograms containing a large variety of diverse particles and corresponding ground-truth segmentation masks (see Section 4).

### Prompt-based particle picking with ProPicker

First, we choose a prompt for the particle of interest. Next, we embed the prompt and segment the tomogram using ProPicker. As tomograms are typically very large, we segment the volume using a strided 3D sliding window approach.

After binarizing the model output ***y*** based on a threshold, we extract predicted particle center coordinates **C** from the resulting segmentation mask using one of two strategies:

- **Clustering-based picking (ProPicker-C)**:We detect clusters by finding connected components [DTT+23]. The centroid of each cluster is a predicted particle center. The precision of this approach can be improved by leveraging prior information about the target particle size by excluding clusters that are significantly smaller or larger than the target particle.
- **Template-matching-based picking (ProPicker-TM)**:We apply a template matching based picker (in this work, we use TomoTwin [Ric+23]) to the input tomogram over regions where our segmentation mask predicts the presence of a particle.

Further information on cluster-based and template-matching-based picking, as well as the hyperparameters of these two strategies, can be found in Appendix B.

Our default variant is ProPicker-C. ProPicker-TM can be used to speed up an existing, well-functioning template matching pipeline. Other use-cases are picking in dense environments where particle clusters are not easily separable (Section 4.1.2), and eliminating false positives from the segmentation mask.

### Particle-specific fine-tuning of ProPicker

Fine-tuning requires (parts of) one or more tomograms with corresponding binary segmentation masks of the particle of interest, as well as a single prompt for the particle of interest. For computational reasons we fine-tune only the segmentation model and keep the prompt encoder frozen and the prompt fixed. Picking with the fine-tuned segmentation model works as in the prompt-based case, i.e. either via clustering or template matching.

Note that the out-of-the-box ProPicker model can help during the annotation of tomograms for fine-tuning, e.g., one can manually select true positives from the picks produced with prompt-based picking.

## 4 Experiments

Before we present our experiments, we describe the model architecture and training dataset ProPicker, and the evaluation setup used throughout all experiments.

**ProPicker Architecture.** We use a slightly modified and larger version of the 3D U-Net used in DeepETPicker [Liu+24] as segmentation model.

As prompt encoder, we use the trained TomoTwin encoder [Ric+23] which maps a 37 × 37 × 37 sub-tomogram (prompt), extracted from one of the tomograms and showing the particle of interest, to a 32-dimensional vector. These embedding vectors are well-suited representations of the particle class of interest, as they have been shown to be useful for particle classification based on cosine similarity [Ric+23].

More details on the U-Net and how we condition it on the encoded prompts are in Appendix A.

**Training Dataset of ProPicker.** We train ProPicker on realistically simulated tomograms from Rice et al. [Ric+23] and Gubins et al. [Gub+20], which have also been used to train TomoTwin. Our training set is a subset of TomoTwin’s training data and consists of 85 tomograms containing a total of 121 unique protein types, as well as gold fiducial markers, vesicles and filaments. Each tomogram contains around 1500 protein instances, each belonging to a set of up to 13 unique protein types. We train on sub-tomograms of size 64 × 64 × 64 extracted from all tomograms. Details on model training are in Appendix C.

**Evaluation Setup.** Following common practice [Moe+21; DTT+23; Ric+23; Liu+24], we measure the particle picking performance with the F1 score of the predicted particle locations (see Appendix D).

Note that to obtain predicted particle locations from raw model outputs, all methods considered in the experiments use classical post-processing routines, e.g., clustering (ProPicker-C) or peak detection (TomoTwin), which rely on particle and dataset-specific hyperparameters that have to be specified by the user. We tune hyperparameters on hold-out calibration data unless explicitly stated otherwise. For more details on hyperparameters and their tuning, see Appendix B.

### 4.1 Prompt-Based Picking

In this Section, we consider prompt-based picking, which requires only a single observation of the particle as prompt, and is, therefore, highly data efficient.

**Baselines.** As baselines, we consider pickers that are able to pick particles based on a single observation or prompt. State-of-the-art methods that fall into this category are TomoTwin [Ric+23] and CryoSAM [Zha+24] introduced in Section 2.

#### 4.1.1 Prompt-Based Picking of Unseen Particles in a Synthetic Tomogram

We first demonstrate that, based on a single prompt, ProPicker can accurately pick particles it has not seen during training in a synthetic tomogram generated with the same simulator as the training data.

**Dataset & Setup.** We consider a regenerated version of the generalization tomogram from Rice et al. [Ric+23; Ric+22], a tomogram with dimensions 200 × 512 × 512 voxels containing 7 unique particles that are not part of our training set.

We applied TomoTwin and ProPicker directly to the tomogram using the same, randomly sampled prompts. Since we were unable to produce any picks with CryoSAM when applying it to the noisy tomogram, we applied it to the clean ground truth tomogram that underlies the simulation of the noisy tomogram to probe the best-case performance. The results for TomoTwin and ProPicker on the original version of the generalization tomogram from the TomoTwin tutorial [Ric+22] are in Appendix E.

Note that for this experiment, we report best-case F1 scores with picking hyperparameters (see Appendix B) tuned directly on the test tomogram. This is because we test on a single tomogram, and to ensure compatibility with the setup in the TomoTwin tutorial [Ric+22], in which the generalization tomogram is used and the hyperparameters are tuned on the test data. We found that the best-case F1 scores are overall very close to those obtained when tuning hyperparameters on hold-out data. All other F1 scores reported in the main paper for all methods were obtained by tuning hyperparameters on hold-out data.

**Results.** As can be seen in Table 1, ProPicker-C performs best on most particles. CryoSAM achieves lower F1 scores than TomoTwin and ProPicker for all particles even though we applied it to clean ground truth tomograms. Upon inspection, we found that CryoSAM’s segmentation masks confuse particles. It is likely that both DINOv2 [Oqu+23] and SAM [Kir+23], which are used in CryoSAM without fine-tuning on cryo-ET data, fail to discriminate between particles even in the absence of noise, highlighting the need for domain-specific training.

**Table 1:** Best-case F1 scores and runtimes (measured on a single NVIDIA L40 GPU) for CryoSAM, TomoTwin, and ProPicker on eight particles not seen during training. The scores for CryoSAM have been obtained on the clean, ground-truth tomogram, as we could not produce good results on the noisy tomogram used as input for TomoTwin and ProPicker.

| Method / PDB | 1avo | 1e9r | 1fpy | 1fzg | 1jz8 | 1oao | 2df7 | Mean | Runtime (s) |
| --- | --- | --- | --- | --- | --- | --- | --- | --- | --- |
| **CryoSAM** | 0.21 | 0.30 | 0.43 | 0.22 | 0.50 | 0.45 | 0.70 | 0.40 | 262 |
| **TomoTwin** (stride=2) | 0.59 | 0.72 | 0.86 | 0.48 | 0.73 | 0.70 | 0.92 | 0.71 | 2620 |
| **TomoTwin** (stride=4) | 0.55 | 0.70 | 0.81 | 0.49 | 0.69 | 0.63 | 0.92 | 0.68 | 362 |
| **TomoTwin** (stride=8) | 0.07 | 0.30 | 0.27 | 0.33 | 0.14 | 0.12 | 0.88 | 0.30 | 80 |
| **ProPicker-TM** | 0.60 | 0.72 | 0.84 | 0.58 | **0.78** | 0.65 | 0.92 | 0.73 | 154 |
| **ProPicker-C** | **0.72** | **0.82** | **0.88** | **0.68** | 0.69 | **0.78** | **0.98** | **0.79** | **77** |

We also compare the runtimes of all methods. In Table 1, we report the time it takes to pick a single particle, not accounting for speedups achievable via caching, e.g., for TomoTwin, the embeddings of sub-tomograms only have to be calculated once if one wants to pick multiple particles within a single tomogram.

TomoTwin is the slowest method as it has to process the tomogram with a very finely-strided (the default stride is 2 voxels) sliding window to achieve high picking performance. ProPicker, too, employs a sliding window approach but due to its 3D segmentation network, we can choose significantly larger strides (our default is 32 voxels). A more extensive analysis of the speed vs. performance trade-off of TomoTwin and ProPicker is in Appendix F.

Our experiment shows that ProPicker can pick unseen particles in a context that is familiar from training with high speed and accuracy.

#### 4.1.2 Prompt-Based Picking in Real-World Tomograms

To further showcase the generalizability of ProPicker, we demonstrate that ProPicker can generalize to real-world tomograms, even though we have trained exclusively on synthetic data. Here, we only compare to TomoTwin which we find to be the stronger baseline, and as we were unable to produce good picks with CryoSAM.

**Dataset & Setup.** We consider the *in-vitro* `DS-10440` dataset by Peck et al. [Pec+24], which contains 7 annotated tomograms. Among other particles, the dataset contains expert annotations for ribosomes and apoferritin. Apoferritin has a molecular weight of 450 kilodalton which is 10.47% of that of the ribosomes in the dataset (4.3 megadalton), and is also smaller in size than the ribosomes.

Our goal is to pick all instances of ribosomes and apoferritin in the tomograms based on a single prompt each. Of the seven available tomograms, we use a single one (`TS 5 4`) to tune the picking hyperparameters, after which we exclude it from the quantitative evaluation of the picking performance.

**Results.** As can be seen in Figure 2 and Figure 3, ProPicker and TomoTwin perform acceptably at picking apoferritin and the ribosomes. ProPicker-TM performs best and ProPicker-C worst. ProPicker-C performs worst due to false positives in the predicted segmentation mask and crowded areas (see, e.g., the zoomed-in region in Figure 2), where several instances of the same particle appear close together, leading to inseparable clusters. This makes cluster-based picking challenging.

**Figure 2:**
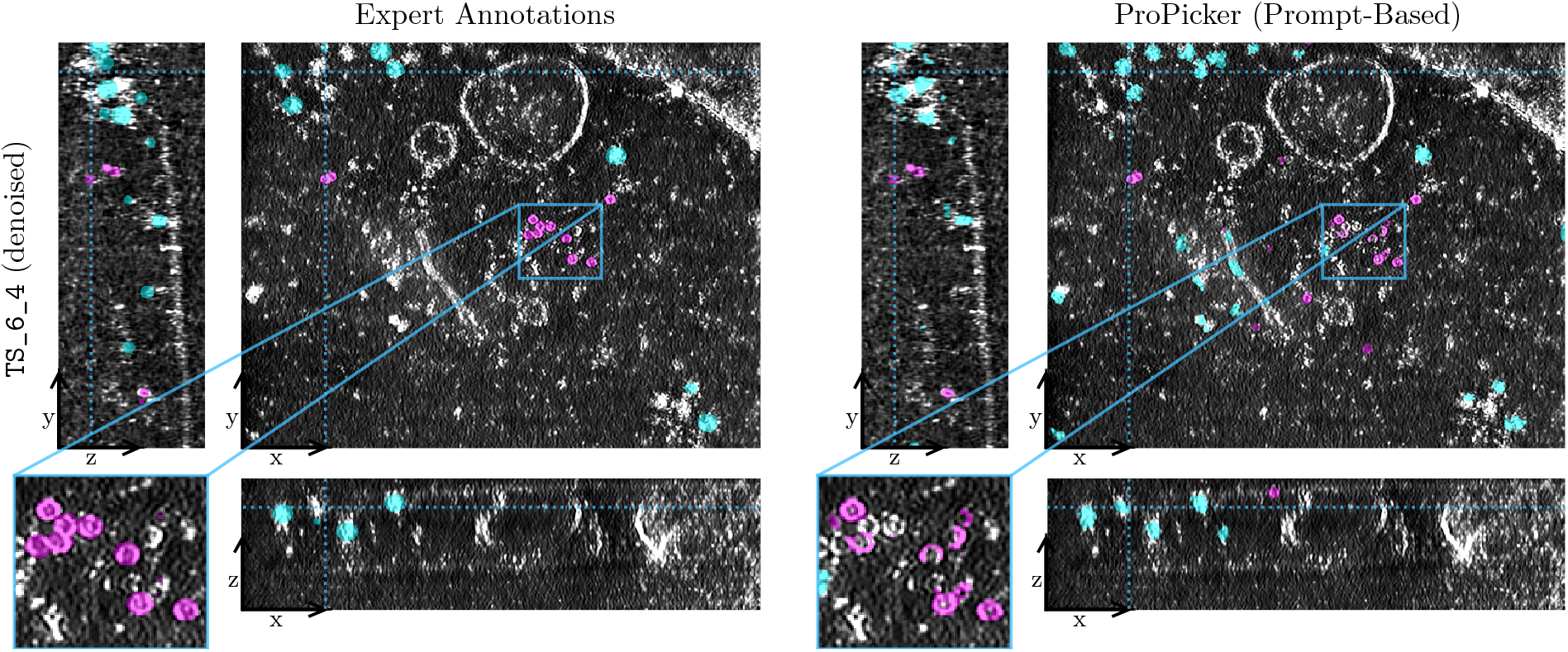
Slices through a single real-world tomogram (TS 6 4) from the DS-10440 dataset [Pec+24] taken from the Cryo ET Data Portal [Erm+24]. Ground-truth annotations and ProPicker segmentation masks for apoferritin and ribosomes are shown in pink and blue respectively. The tomogram underlying the segmentation masks has been denoised by Peck et al. [Pec+24]. The ProPicker masks have been obtained from the raw, noisy tomogram (not shown). The images are high resolution, and we recommend readers to zoom in.

**Figure 3:**
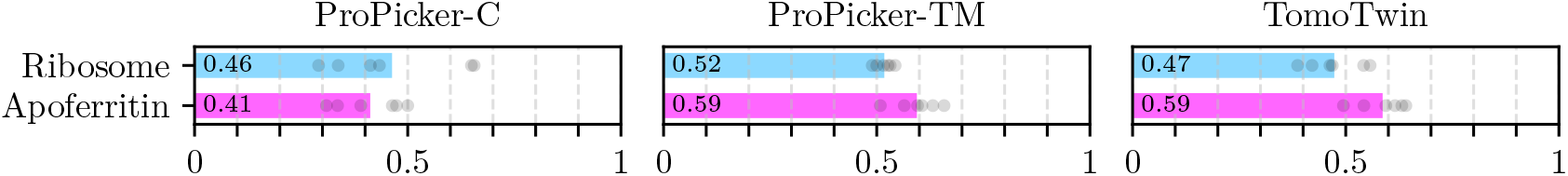
Average F1 scores for apoferritin and ribosomes picked in 6 raw real-world tomograms from the DS-10440 dataset [Pec+24] (excluding TS 5 4). The gray points show the scores for the 6 individual tomograms. For ProPicker-TM, we used TomoTwin as template matching picker

ProPicker-TM and TomoTwin achieve better picking performances. Overall, ProPicker-TM performs on par (apoferritin) or better (ribosomes) than TomoTwin on its own and at the same time offers large speedups: On average, the ProPicker segmentation masks used for ProPicker-TM masked out 99.5% of all voxels for apoferritin and 97.3% of all voxels for ribosomes, resulting in relative speedups over TomoTwin which are similar to those shown in Table 1.

We could not produce good picks for the other particles contained in the tomogram, which are virus-like particles, beta amylase, beta galactosidase, and thyroglobulin, with neither TomoTwin, nor ProPicker. Beta amylase, beta galactosidase, and thyroglobulin are very small, produce little contrast and are therefore considered very hard to pick [Pec+24].

### 4.2 Improving Over Prompt-Based Picking with Fine-Tuning

In some cases, ProPicker is able to accurately pick particles based on a single prompt, but there are particles which neither ProPicker nor TomoTwin are able to locate. Here, we show that in such cases ProPicker’s performance can be improved through fine-tuning on little data, and that fine-tuning ProPicker can be more data-efficient than training a state-of-the-art picker from scratch.

**Baselines.** Aside from TomoTwin, we compare to particle-specific deep-learning-based pickers, specifically to the recent DeepETPicker method [Liu+24]. DeepETPicker is segmentation-based and uses a convolutional 3D U-Net architecture. It is a suitable baseline due to its state-of-the-art performance and data-efficiency [Liu+24]. We also tested DeepFinder [Moe+21], which is conceptually similar to DeepETPicker, but found that its performance and data efficiency are lower than those of DeepETPicker, which is in line with the comparison of the two methods by Liu et al. [Liu+24].

We consider two variants of DeepETPicker, which we train as segmentation models, i.e., with particle segmentation masks as targets. For the first variant, “DeepETPicker (1 Class)”, we train one DeepETPicker model for each particle separately. This is the same setup as when we fine-tune ProPicker. For the second variant, “DeepETPicker (8 Classes)”, we train one DeepETPicker model to pick all 8 particles *simultaneously*. Moebel et al. [Moe+21] observed that multi-class training can yield substantial improvements in performance for difficult-to-pick particles. In contrast to the single class setup, the multi-class setup requires annotations for *all 8 particles*.

Finally, recall that ProPicker and DeepETPicker use a similar 3D U-Net architecture, but ProPicker’s model has around 42× more parameters. To ensure that the difference in model size does not distort our experiments, we re-ran parts of the experiments with a variant of DeepETPicker in which we scaled its U-Net to the same size as ProPicker’s U-Net (100 million parameters). The average performance of the larger U-Net is very similar to that of the original and smaller one. Details on this, and a comparison of runtimes for training/fine-tuning and inference with ProPicker and DeepETPicker are in Appendix G).

#### 4.2.1 Prompt-Based Picking and Fine-Tuning for Unseen Particles in Synthetic Tomograms

We first fine-tune ProPicker on a set of synthetic tomograms and show that, depending on the particle, fine-tuning ProPicker can be more data-efficient than training DeepETPicker from scratch.

**Dataset & Setup.** We consider a set of 8 tomograms from TomoTwin’s training set, each of which contains instances of 8 unique particles. For our fine-tuning experiment, we first re-trained both TomoTwin and ProPicker, exlcuding the 8 tomograms containing the 8 particles from the training set. As, by design, TomoTwin’s original training set contains particles whose 3D structures are all very different from one another (see [Ric+23] for details), all 8 particles are hard, unseen examples for ProPicker and TomoTwin, but the context is similar to that during training as the tomograms come from the same simulator.

Out of the 8 available tomograms with shape 200 × 512 × 512, we used up to 4 for training/fine-tuning, one for validation and for tuning picking hyperparameters, and the remaining 3 for testing.

**Results.** Figure 4 shows the results of our experiment. Both TomoTwin and prompt-based picking with ProPicker-C yield satisfactory or even near-perfect (`6mrc`) scores for some particles, while for other particles, the performance is poor.

**Figure 4:**
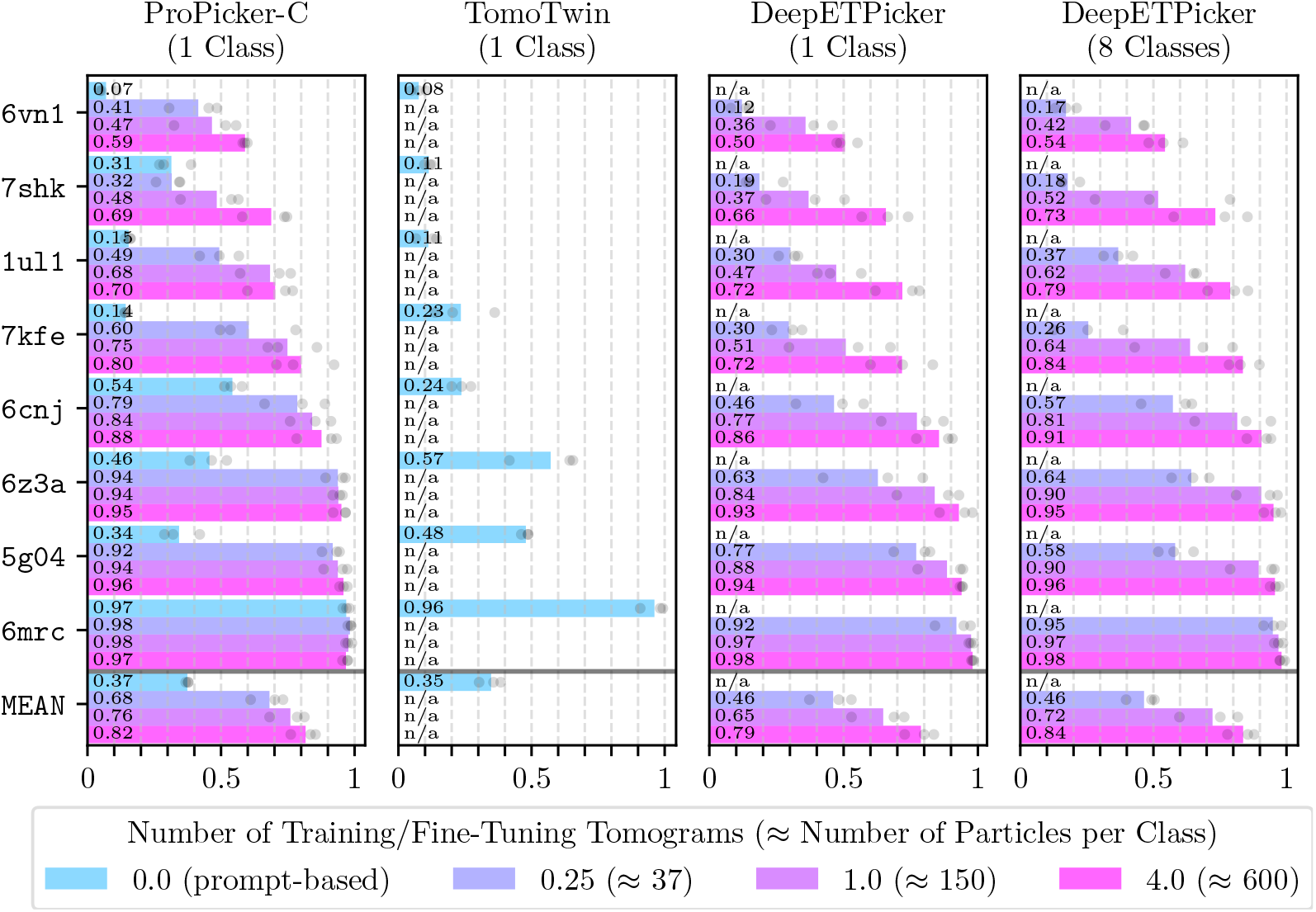
Average F1 scores of ProPicker-C, TomoTwin and DeepETPicker versus amount of training/fine-tuning data across 3 test tomograms. The gray points show the scores for the 3 individual test tomograms.

**Figure 5:**
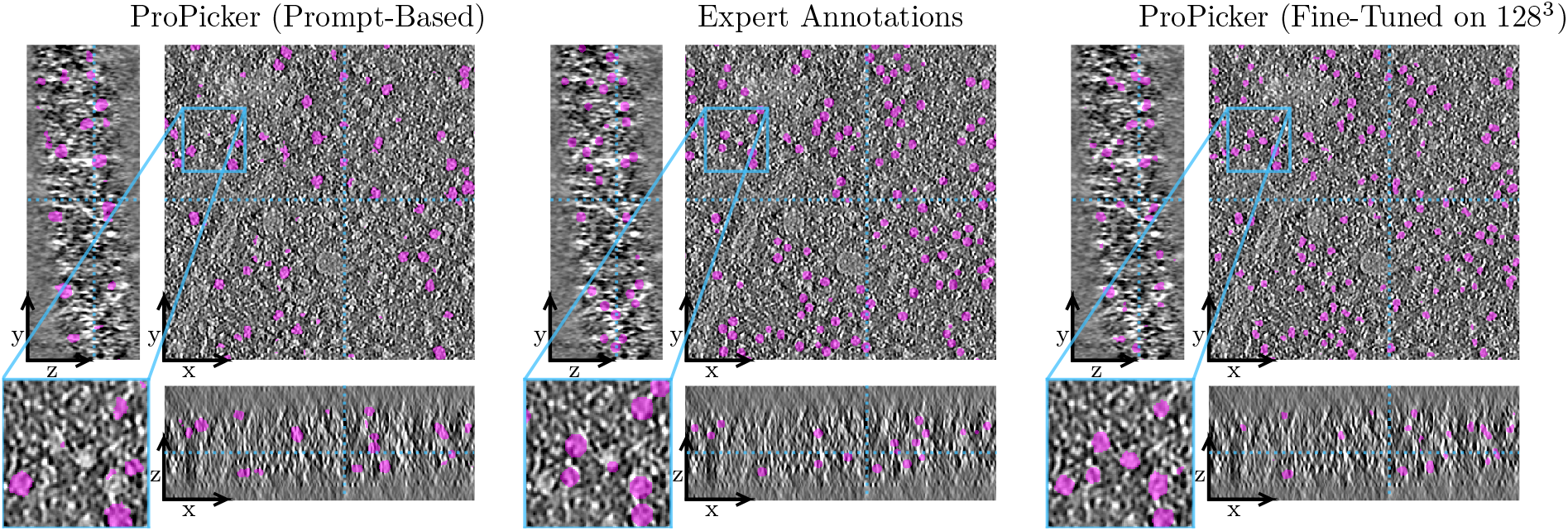
Slices through a single real-world tomogram (TS 030) from EMPIAR-10988 and expert annotations by De Teresa-Trueba et al. [DTT+23]. Fine-tuning ProPicker on a sub-tomogram of shape 128 × 128 × 128 (right panel) significantly increases the number of picked particles compared to prompt-based picking (left panel). The images are high resolution, and we recommend readers to zoom in.

Fine-tuning ProPicker boosts the picking performance: Depending on the particle, fine-tuning on as little as 25% of a single tomogram improves the performance up to 4× (`7kfe`). The performance of the ProPicker models saturates quickly as more data is used for fine-tuning.

If little data *or* only annotations for the particle of interest are available, fine-tuning ProPicker yields superior performance compared to both variants of DeepETPicker: When training/fine-tuning on up to one single tomogram, the fine-tuned ProPicker-C outperforms DeepETPicker (1 Class) and DeepETPicker (8 Classes). The performance gap between all methods narrows as more data is used.

DeepETPicker (8 classes) performs on par or slightly better than the best ProPicker-C pickers if enough training data (4 tomograms) is available. The improved performance over the single-class version comes at the cost of having to annotate *all* particles even if one is only interested in a single one.

#### 4.2.2 Prompt-Based Picking and Fine-Tuning for Ribosomes in *In-Situ* Tomograms

Our final experiment is on a set tomograms showing crowded sections of *S. pombe* cells. Such *in-situ* data is among the most interesting, yet most challenging types of data that is studied with cryo-ET.

**Dataset & Setup.** We consider tomograms from the `EMPIAR-10988` dataset by De Teresa-Trueba et al. [DTT+23], which mainly show ribosomes within *S. pombe* cells.

Our goal is to pick all ribosomes in 8 (defocus-only) tomograms. We fine-tune ProPicker on center-crops of different sizes which we extract from a single tomogram (`TS 029`). We use center-crops from another tomogram (`TS 030`) as validation data, to choose a prompt for TomoTwin and ProPicker, and for tuning picking hyperparameters. As a reference, we train DeepETPicker from scratch with the same setup. More details on model training and evaluation in this experiment are in Appendix H.

**Results.** In Figure 6, we see that TomoTwin outperforms ProPicker-C in prompt-based picking, but fine-tuning even on the smallest sub-tomogram, which contains only 65 particles, results in almost 50% improvement over TomoTwin. Due to its pre-training, the fine-tuned ProPicker-C outperforms DeepETPicker in the low data regime, but the difference in performance diminishes as more data becomes available.

**Figure 6:**
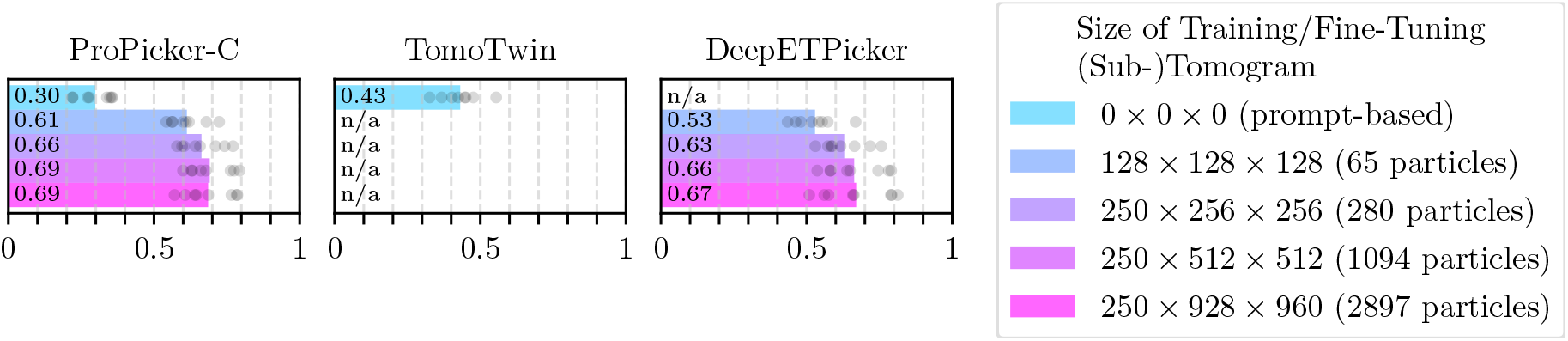
F1 scores of ProPicker-C, TomoTwin and DeepETPicker versus the amount of training/fine-tuning data for the ribosomes in the EMPIAR-10988 tomograms. The colored bars represent the average scores across 8 (defocus-only) tomograms (excluding TS 029 and TS 030), and the gray points show the scores for the individual tomograms.

In addition to annotations for ribosomes, the `EMPIAR-10988` dataset also contains annotations for fatty acid synthases (FAS). We were unable to produce good prompt-based picking results for FAS with ProPicker and TomoTwin, and, therefore, we do not include them in the fine-tuning experiment. The poor performance is not surprising, as FAS is difficult to pick, and even pickers specifically trained on `EMPIAR-10988` for picking FAS achieve low F1 scores [DTT+23].

## 5 Discussion

In this work, we have proposed ProPicker, a particle picking method for cryo-ET that uses a promptable 3D segmentation model trained on a large and diverse synthetic dataset. ProPicker can target any particle of interest by specifying a single example (prompt), and achieves state-of-the-art performance or nearly state-of-the-art performance while being up to an order of magnitude faster. Moreover, fine-tuning ProPicker outperforms state-of-the-art particle-specific pickers if only limited training data is available, which is often the case in practice.

ProPicker has the flexibility of pickers like TomoTwin [Ric+23] and CryoSAM [Zha+24]. At the same time, ProPicker is easily finetunable and thus achieves the better performance of methods like DeepETPicker [Liu+24] which are trained from scratch for each specific dataset or particle of interest. Compared to DeepETPicker, ProPicker is more data-efficient, i.e., requires less training/fine-tuning data due to its pre-training.

### Limitations

Limitations of ProPicker compared to existing particle pickers are the following:

First, the performance of prompt-based picking with ProPicker depends on the choice of the prompt. However, fluctuations are small for most particles, especially for such for which prompt-based picking works well overall (see Appendix I). This limitation is not unique to ProPicker and also applies to all flexible pickers that use an example or prompt. We recommend trying a few prompts for each particle. Developing principled ways to find prompts that yield high performance is an interesting direction for future research.

Second, we found that prompt-based picking performance of ProPicker-C to be worse and less robust compared to TomoTwin in some cases (see results in Section 4.1.2 and Section 4.2.2). In such cases, the ProPicker-TM variant with TomoTwin template matching can be a viable option since it can reach the same or even higher performance (see Section 4.1.2) while at the same time providing significant speedups.

A limitation regrading fine-tuning ProPicker is that its large model size results in longer training and inference times compared to from-scratch training with DeepETPicker (see Appendix G.2). However, the higher data efficiency of fine-tuning ProPicker compensates for that in practice, as annotation of training data can take weeks [Pec+24].

### Outlook

An ideal particle picker can accurately detect any particle in any tomogram based on a single prompt or instruction, with high speed and minimal human intervention. Such a picker would significantly facilitate and accelerate cryo-ET research, where a major bottleneck is particle picking [Pec+24; MS25].

While initial results toward a fast, universal picker achieved with, e.g., TomoTwin and ProPicker are promising, existing methods are not yet truly universal [HZB24; MS25] and struggle with some particles and contexts. More work is needed to improve their performance and robustness.

We believe that a promising next step towards a robust universal picker is to collect large and diverse datasets of real-world tomograms with ground-truth particle annotations for a variety of particles for training, as it is widely accepted that training such datasets improves the robustness of deep learning models [Rad+21; Fan+22; LH24]. Large-scale efforts to collect such datasets have already been initiated [Ish+23; Erm+24], and methods like ours can help researchers to reduce the manual annoation effort needed for this task. For example, a promising idea is to use an iterative approach where trained models help annotating data that is then used to re-train the models. Such an iterative approach has already been applied in the cryo-ET domain for the training of membrane segmentation models [Lam+22; Lam+24].

While more and better training data will improve models, the high variability of biological macro-molecules and their cellular contexts might place a truly universal picker that can robustly generalize to arbitrary particles and contexts out of reach. Therefore, we see the most promising direction in methods that combine diverse pre-training with task-specific fine-tuning, as they enable a data-efficient workflow to obtain a high-performance picker.

## Data Availability

Simulated tomograms used to train ProPicker, and for the fine-tuning experiments in Section 4.2.1 are part of TomoTwin’s training and validation data, which can be downloaded here: https://doi.org/10.5281/zenodo.6637456.

We also used tomograms from the SHREC 2021 dataset for training. The SHREC 2021 dataset can be downloaded here: https://doi.org/10.34894/XRTJMA.

The `DS-10440` dataset used for the experiments in Section 4.1.2 can be downloaded from the CryoET Data Portal: https://cryoetdataportal.czscience.com/datasets/10440.

The `EMPIAR-10988` datasets used for the experiments in Section 4.2.2 can be downloaded here: https://doi.org/10.6019/EMPIAR-10988.

## Code Availability

Code for prompt-based picking, as well as training and fine-tuning ProPicker is available on GitHub:https://github.com/MLI-lab/ProPicker.

## Author Contributions

S.W., Z.F., M.S., and R.H. conceptualized the project and planned the experiments. S.W. and Z.F. developed the code and conducted the experiments. S.W. performed the data analysis and prepared the illustrations. S.W. and Z.F. wrote the initial draft, which was reviewed and edited by all authors. M.S. and R.H. supervised the project and acquired funding.

## Appendix

### A Details on the Promptable Segmentation Model Architecture

Here, we provide details on our concrete choice for the segmentation model and how we condition it on a prompt, i.e., an observation of the particle of interest.

#### A.1 Segmentation Model Architecture

We use a well-established convolutional 3D U-Net [RFB15], which is an encoder-decoder architecture (see Figure 1), as our segmentation model. The U-Net’s architecture follows the one used in DeepETPicker [Liu+24], but we scaled it up for better performance on the large pre-training dataset: The encoder consists of 5 convolution-based spatial downsampling layers, and the corresponding decoder has 5 convolution-based spatial upsampling layers. The layers process 3D volumes with 108, 216, 432, and 864 channels, respectively, which amounts to 100 million trainable parameters. For comparison: the default U-Net in DeepETPicker, too, uses 5 up- and downsampling layers but with 24, 48, 96 and 192 channels, which results in 2.4 million trainable parameters.

Another difference between the U-Net we use in ProPicker and the one used in DeepETPicker is that we use instance normalization [UVL16] instead of batch normalization [IS15] as the former generalizes better and allows smaller batches during training, which is favorable for large models like ours.

#### A.2 Conditioning the Segmentation Model on the Prompt

We condition each of the decoder’s 5 spatial upsampling layers on the encoded prompt *ε* (***p***) *∈* ℝ_*d*_ with the FiLM [Per+18] approach, which works as follows: Let *c* be the number of channels (features) of an intermediate 3D feature map after upsampling. First, we multiply the encoded prompt *ε* (***p***) *∈* ℝ_*d*_ with two (learnable) matrices ***A, B*** *∈* ℝ_*c*×*d*_. We then map each channel *k ∈ {*1, …, *c}*, with an affine transformation with slope (***A****ε* (***p***))_k_*∈* ℝ and intercept ***(B****ε* (***p***))_*k*_ *∈* ℝ, which gives the conditioned feature map. We use a separate pair of learnable matrices (***A, B***) for each of the 5 up-sampling layers.

### B Picking Hyperparameters

All methods we consider in this work do not directly predict particle positions. Instead, they produce 3D segmentation masks (ProPicker, CryoSAM, DeepETPicker) or heatmaps (TomoTwin) from which particle locations are derived using clustering (ProPicker-C, CryoSAM, DeepETPicker) or peak-detection (ProPicker-TM, TomoTwin).

Both clustering and peak-detection use particle-specific hyperparameters that have to be determined by the user, usually using a small annotated calibration dataset. Here, we detail the extraction of particle coordinates from segmentation mask or heatmap outputs of each method considered in the paper:

- **TomoTwin**:TomoTwin’s peak-detection mainly relies on four thresholds: A minimum value for when a candidate point is considered a peak, a threshold for flood-filling which is used to enlarge the peak and two thresholds for the minimum and maximum number of voxels the flood-filled peak must consist of. The TomoTwin software package has an option to tune the peak threshold and the two size thresholds, which we used in all experiments. For all experiments on synthetic data and on `DS-10440` [Pec+24], we found the default flood-filling threshold (`findmax_tolerance`) of 0.2 to yield good performance. For the experiments involving the EMPIAR 10988 dataset (Section 4.2.2), the default threshold resulted in poor performance, so we had to manually tune it, and found 0.015 to be optimal.
- **ProPicker-C**:The output of ProPicker’s segmentation model is a sigmoided 3D volume ***y*** *∈* [0, 1]^*n*×*n*×*n*^ of the same shape as the input tomogram. To produce particle positions via clustering, we first binarize the sigmoided volume based on a threshold, then cluster the binary volume based on a connected component analysis (which is inspired by DeePiCt [DTT+23]), and, finally, remove clusters that are too large or too small based on a threshold. The maximum and minimum acceptable cluster sizes can be chosen relative to the particle size. We optimize the binarization threshold and the two size thresholds using a search on labeled calibration data. An example for how tuning each hyperparameter affects the performance of ProPicker-C is shown in Figure A1.
- **ProPicker-TM**:ProPicker-TM applies template matching - we use TomoTwin for that - to a segmentation mask produced with ProPicker. This requires first binarizing the sigmoided ProPicker output. In all experiments, we used 0.1 as threshold. Then we applied TomoTwin, ignoring all locations where the mask is 0, which is natively supported by TomoTwin. The final particle coordinates are, therefore, produced with TomoTwin’s peak-detection routine with hyperparameters as discussed above.
- **CryoSA:**CryoSAM outputs a binary segmentation mask. The implementation by Zhao et al. [Zha+24] does not contain a method to produce particle locations based on this mask, so we applied the same clustering as for ProPicker-C. As CryoSAM’s output is already binary, we only optimized the cluster size thresholds.
- **DeepETPicker**:The implementation of DeepETPicker provided by Liu et al. [Liu+24] comes with a cluster-based picking routine. However, we found that the clustering method used for ProPicker-C yields better results for DeepETPicker on synthetic data. Therefore, and because it makes the comparison more principled, we used the same clustering routine as for ProPicker-C instead of DeepETPicker’s and optimized the same hyperparameters.

**Figure A1:**
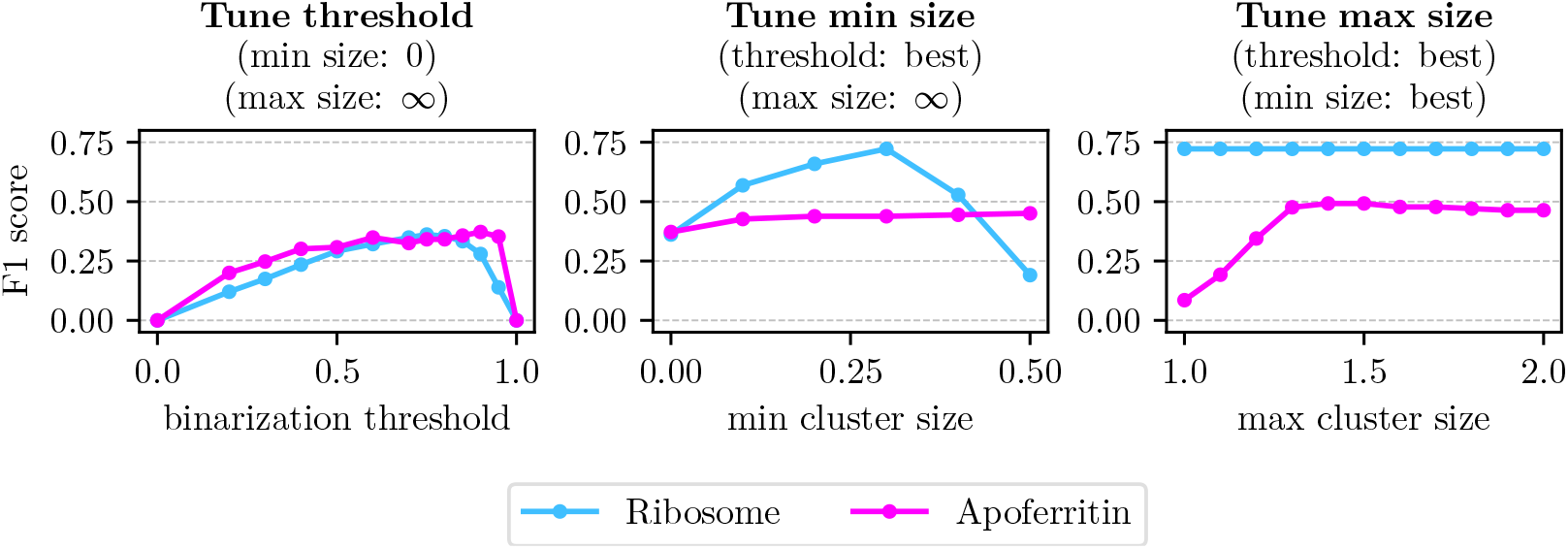
Picking F1 score versus the three hyper-parameters of ProPicker-C’s cluster-based picking routine for a single tomogram from the setup of Section 4.1.2. The figure illustrates the process of first tuning the binarization threshold (left panel), then using the optimal binarization threshold to tune the minimum cluster size (center panel), and finally tuning the maximum cluster size based on the optimal binarization threshold and minium cluster size found before. The cluster sizes on the x-axes are relative values, defined as the number of positive voxels in the cluster divided by the estimated size of the particle.

### C Training ProPicker

To reduce the computational cost of training, we keep the TomoTwin prompt encoder frozen during training (and fine-tuning). We train the segmentation model using the Adam optimizer [KB15] with a fixed learning rate of 0.01.

For each gradient step with Adam, we first randomly sample a batch of 8 sub-tomograms. Each sub-tomogram can contain particles that belong to up to 13 unique classes (see Section 4). Next, we randomly sample 8 prompts. Each prompt corresponds to one particle class of which instances may be contained in the sub-tomogram. We pass each sub-tomogram and its corresponding 8 prompts through the conditional segmentation model. This yields a total of 64 = 8 *·* 8 predicted segmentation masks. Finally, we compute the average voxel-wise binary cross-entropy between the model outputs and the 64 single-class target masks as loss for the calculation of gradients.

### D Picking F1 Score

We evaluate the picking performance of all methods using the F1 score, which is defined as the harmonic mean of precision and recall. Computing the F1 score requires a criterion based on which we consider a predicted particle position (given by 3 coordinates) to be a true-positive.

As our main baseline in this work is TomoTwin, we adopt the criterion used by Rice et al. [Ric+23], who consider a predicted position a true-positive if the intersection-over-union (IoU) between a cubic box centered at the predicted position and a cubic box of the same size centered at a ground-truth particle position is at least 0.6. The side length of the cubic box is chosen according to the size of the particle of interest.

### E Performance of ProPicker and TomoTwin on the Original Generalization Tomogram from the TomoTwin Paper

**Table A1:** Best-case picking F1 scores for TomoTwin and ProPicker on the original version of the generalization tomogram that is used in the official TomoTwin tutorial [Ric+22].

| Method / PDB | 1avo | 1e9r | 1fpy | 1fzg | 1jz8 | 1oao | 2df7 | Mean | Median |
| --- | --- | --- | --- | --- | --- | --- | --- | --- | --- |
| **TomoTwin** (stride=2) | 0.71 | 0.80 | **0.70** | **0.90** | 0.85 | 0.86 | **0.99** | **0.83** | 0.84 |
| **ProPicker-TM** | 0.68 | 0.78 | 0.51 | 0.88 | **0.89** | **0.93** | 0.98 | 0.81 | 0.84 |
| **ProPicker-C** | **0.87** | **0.91** | 0.33 | 0.85 | 0.70 | 0.91 | 0.97 | 0.79 | **0.86** |

### F Picking Speed

Both TomoTwin and ProPicker process the tomogram in a 3D sliding window fashion. Therefore, inference time is cubically related to the window stride. However, large strides (small overlap) often results in low detection performance. Here, we explore this trade-off. To quantify speed, we report the throughput in tomograms per GPU hour measured on a single NVIDIA L40 GPU for picking a single particle of interest in a tomogram of size 200 × 512 × 512.

As the speed at which a particle can be reliably picked depends on, e.g., the particle’s size (especially for template matching methods like TomoTwin), we measure picking speed on a set of 11 tomograms which contain 108 unique particle types in total. Both TomoTwin and ProPicker have seen all of these particle types during training, but within different tomograms.

**Figure A2:**
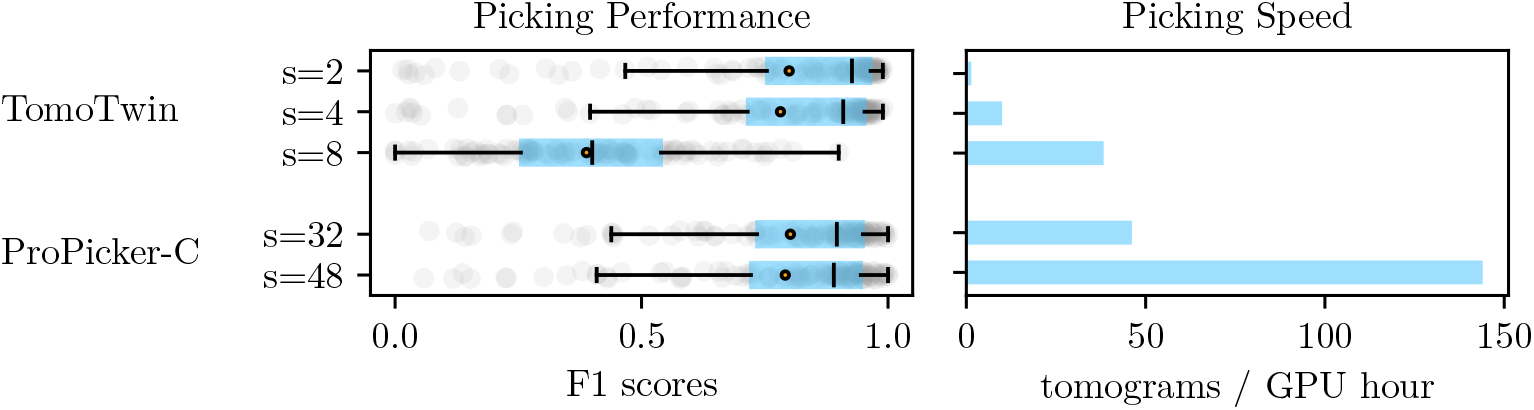
Best-case F1 scores and speed for ProPicker-C and TomoTwin for particles seen during training. In the boxplots, vertical markers are medians, and circles are means.

As can be seen in Figure A2, TomoTwin with *s* = 2 performs best but is the slowest (*<* 2 tomograms / GPU hour). Increasing the stride to *s* = 4 results in a small drop in performance (*™*0.03 mean and median F1 score) and a large relative speedup of around 8×. ProPicker-C with *s* = 32 performs as well as TomoTwin for *s* = 4, but is 4.5× faster.

Achieving a picking speed similar to that of ProPicker-C with TomoTwin by increasing the stride to *s* = 8 comes at the cost of a large drop in performance. ProPicker-C on the other hand can be sped up further: Increasing the stride from *s* = 32 to *s* = 48 increases the speed by a factor 3.3 (15× speedup compared to TomoTwin with *s* = 4) while achieving almost the same performance.

ProPicker’s speed advantage over TomoTwin, and template matching in general, is due to its use of a segmentation network for picking rather than the template matching approach of comparing candidate particles to a template on a fixed grid. The reason is that the “resolution” at which template matching can pick particles depends directly on the granularity of the grid. In general, small strides (fine grids) are required to obtain high accuracy. In contrast, segmentation-based picking does not rely on grid resolution in the same way. Here, smaller strides (increased sub-tomogram overlap) instead improve the consistency of the full tomogram segmentation mask during inference with the sliding window approach.

Other picking methods that use convolutional models for segmentation like DeepFinder [Moe+21], DeePiCt [DTT+23] or DeepETPicker [Liu+24] offer favorable speed-performance trade-offs similar to ProPicker, but are strictly limited to picking particles seen during training.

### G Influence on ProPicker’s Model Size on Performance and Runtime vs. DeepETPicker

While the architectural difference between ProPicker’s and DeepETPicker’s 3D U-Nets are minor, ProPicker’s U-Net is significantly larger to cope with pre-training on a large dataset. ProPicker’s U-Net has 100 million trainable parameters and is, therefore, around 41× larger than DeepETPicker’s default U-Net. Here, we compare the impact of the larger model size on picking performance and runtimes for training and inference.

#### G.1 Influence of the Model Size on the Performance

To ensure a fair comparsion of the picking performance of DeepETPicker and ProPicker, we compared the performance of the original DeepETPicker U-Net with 2.4 million parameters to a scaled-up version with 100 million parameters, which is very similar to the ProPicker U-Net.

We re-ran 6 trainings from the experiment discussed in Section 4.2.1, where we trained DeepETPicker for individual particles on synthetic tomograms. Specifically, we trained the 100 million parameter DeepETPicker U-Net from scratch to pick `1ul1` and `7shk` particles. Like in Section 4.2.1, we trained a separate model for each particle on 25% of a tomogram, a single tomogram, and 4 tomograms, respectively.

For `1ul1`, we obtained mean F1 scores across the 3 test tomograms of 0.27, 0.53 and 0.72 for the large U-Net trained when training on 0.25, 1 and 4 tomograms, respectively. The corresponding values for the small U-Net reported in Figure 4 are 0.30, 0.47 and 0.72.

For `7shk`, the large U-Net achieves mean F1 scores of 0.12, 0.43 and 0.66, which are worse or on-par compared to the scores of the small U-Net reported in Figure 4, which are 0.19, 0.37, 0.66.

The average F1 score obtained by the large U-Net across the 6 training runs (0.46) is very close to the average F1 score obtained by the small U-Net (0.45).

#### G.2 Influence of the Model Size on Runtimes

In the setup of Section 4.2.1, inference with DeepETPicker takes 17s vs. 85s with ProPicker. Training DeepETPicker on 1 tomogram until convergence takes 1:45h, fine-tuning ProPicker on the same data takes 4:45h. We measured all runtimes on a single NVIDIA-L40 GPU. Both methods provide fast enough inference for efficient analysis, especially compared to the hour-long runtimes of widely used template matching methods.

However, we emphasize that ProPicker requires less annotated training data than DeepETPicker. This outweighs slightly slower training and inference, as the (manual) annotation of tomograms for training/fine-tuning data is the major bottleneck in cryo-ET and can take weeks [Pec+24].

### H Technical Remarks on the Experiment in Section 4.2.2

This appendix contains two remarks on the fine-tuning experiment discussed in Section 4.2.2.

#### H.1 Remark on Evaluation Metrics

The F1 scores shown in Figure 6 are lower than the ones reported by De Teresa-Trueba et al. [DTT+23] for DeepFinder, even though DeepETPicker generally outperforms DeepFinder [Liu+24]. This is because we use a stricter true-positive criterion (see Appendix D) in order to be consistent with the other experiments in the paper and because we only used (parts of) a single tomogram for training.

#### H.2 Remark on the Choice of the Loss Function

We obtained all results for training/fine-tuning DeepETPicker and ProPicker discussed in Section 4.2.2 using the binary cross-entropy loss and binary particle segmentations produced by De Teresa-Trueba et al. [DTT+23] as targets. Binary cross-entropy is ProPicker’s default training and fine-tuning loss, whereas DeepETPicker uses the Dice loss [Sud+17] as default. When using the Dice loss, we observed overlapping segmentation masks for individual particles due to the highly crowded nature of the *S. pombe* cells. Such overlapping parts cannot be easily separated via clustering and, therefore, yield lower picking performance: The F1 score when training DeepETPicker on the full tomogram with the Dice loss was around 0.1 lower.

### I Dependence of Picking Performance on the Prompt

For prompt-based picking with ProPicker, the user has to manually extract a prompt from a tomogram at inference. As the tomogram typically contains many instances of the particle of interest, a natural question is how the performance of ProPicker depends on the concrete choice of the prompt.

We investigate this prompt-dependence on three synthetic tomograms, each containing 10 unique particles which are all contained in the training set. We randomly sampled 10 prompts for each particle and evaluated the best-case picking F1 score. Results are shown in Figure A3.

**Figure A3:**
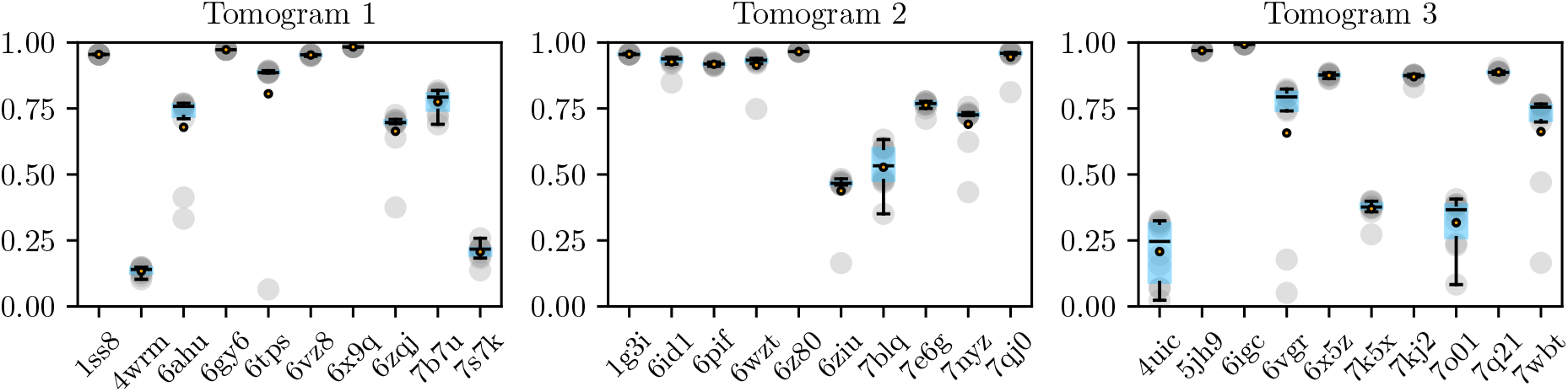
Dependence of the (best-case) picking F1 score of ProPicker-C on the choice of the prompt for two synthetic tomograms containing particles which were seen during training. In the boxplots, vertical markers are medians, and circles are means.

**Figure A4:**
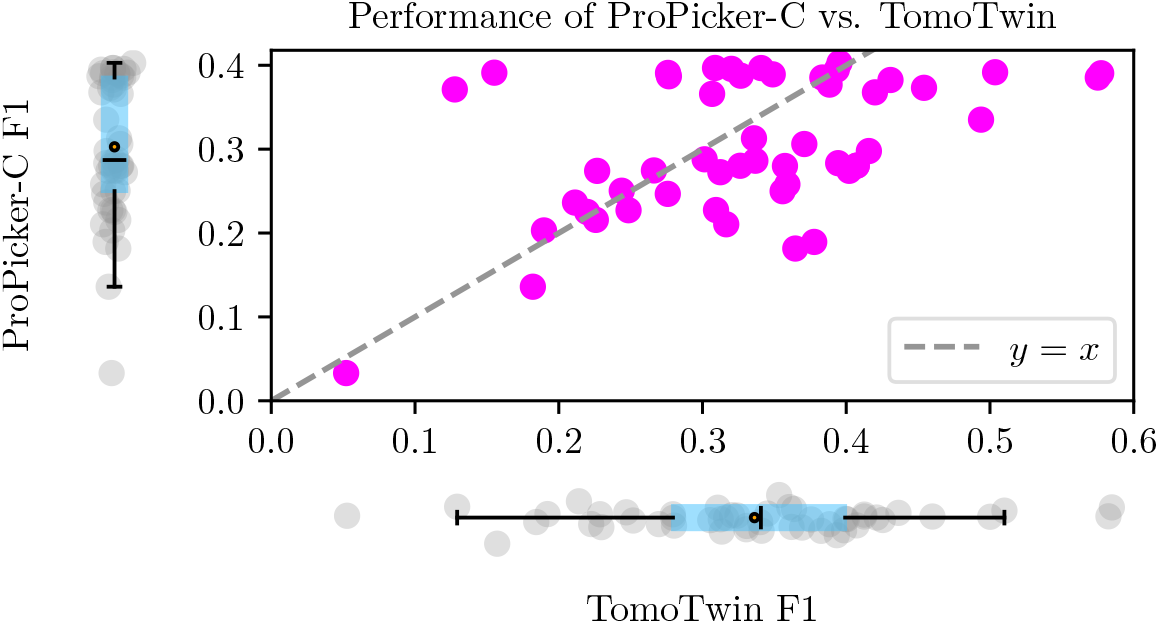
Picking performance of ProPicker-C vs. that of TomoTwin one tomogram (TS 030) from the EMPIAR-10988 dataset for 50 randomly sampled prompts. We optimized picking hyperparameters (see Appendix B) for each prompt separately. In the boxplots, vertical markers are medians, and circles are means.

We performed a similar experiment for the setup of picking ribosomes in the *S. pombe* cells (`EMPIAR-10988`) as discussed in Section 4.2.2. Here, we used 50 random prompts and also evaluate TomoTwin’s performance. Results are shown in Figure A4.

As can be seen in Figure A3 and Figure A4, the picking performance of ProPicker-C does vary with the choice of the prompt. Variations are negligible for particles for which prompt-based picking works well overall. For particles for which the overall performance is lower, e.g. `7blq` (Tomogram 2), `4uic` (Tomogram 3) or the ribosomes in the *S. pombe* cells (Figure A4), the variance is larger.

TomoTwin’s performance, too, depends on the choice of the prompt, as can be seen in Figure A4. We also find that the performance of TomoTwin and that of ProPicker-C are moderately correlated (Pearson correlation coefficient *r*_*xy*_ = 0.49). This is likely because ProPicker uses TomoTwin as prompt encoder.

## References

[Bep+19] T. Bepler, A. Morin, M. Rapp, J. Brasch, L. Shapiro, A. J. Noble, and B. Berger. “Positive-unlabeled convolutional neural networks for particle picking in cryo-electron micrographs”. In: Nature Methods 16.11 (2019).

[Bod+23] S. Bodakuntla, C. C. Kuhn, C. Biertümpfel, and N. Mizuno. “Cryo-electron microscopy in the fight against COVID-19—mechanism of virus entry”. In: Frontiers in Molecular Biosciences 10 (2023).

[Boh+00] J. Bohm, A. S. Frangakis, R. Hegerl, S. Nickell, D. Typke, and W. Baumeister. “Toward detecting and identifying macromolecules in a cellular context: template matching applied to electron tomograms”. In: Proceedings of the National Academy of Sciences 97 (2000).

[CL+24] S. Cruz-León et al. “High-confidence 3D template matching for cryo-electron tomography”.In: Nature Communications 15.1 (2024).

[DTT+23] I. De Teresa-Trueba et al. “Convolutional networks for supervised mining of molecular patterns within cellular context”. In: Nature Methods 20.2 (2023).

[Erm+24] U. Ermel, A. Cheng, J. X. Ni, J. Gadling, M. Venkatakrishnan, K. Evans, J. Asuncion, A. Sweet, J. Pourroy, Z. S. Wang, et al. “A data portal for providing standardized annotations for cryo-electron tomography”. In: Nature Methods 21.12 (2024).

[Fan+22] A. Fang, G. Ilharco, M. Wortsman, Y. Wan, V. Shankar, A. Dave, and L. Schmidt. “Data determines distributional robustness in contrastive language image pre-training (clip)”. International Conference on Machine Learning (2022).

[Gen+23] E. Genthe, S. Miletic, I. Tekkali, R. Hennell J., T. C. Marlovits, and P. Heuser. “PickYOLO: Fast deep learning particle detector for annotation of cryo electron tomograms”. In: Journal of Structural Biology 215.3 (2023).

[Gub+20] I. Gubins, M. L. Chaillet, G. van Der Schot, R. C. Veltkamp, F. Förster, Y. Hao, X. Wan, X. Cui, F. Zhang, E. Moebel, et al. “SHREC 2020: Classification in cryo-electron tomograms”. In: Computers & Graphics 91 (2020).

[HZB24] Q. Huang, Y. Zhou, and A. Bartesaghi. “MiLoPYP: self-supervised molecular pattern mining and particle localization in situ”. In: Nature Methods 21.10 (2024).

[HS21] R. K. Hylton and M. T. Swulius. “Challenges and triumphs in cryo-electron tomography”. In:iScience 24.9 (2021).

[IS15] S. Ioffe and C. Szegedy. “Batch normalization: accelerating deep network training by reducing internal covariate shift”. International Conference on Machine Learning (2015).

[Ish+23] A. Ishemgulova, A. J. Noble, T. Bepler, and A. D. Marco. Preparation of labeled cryo-ET datasets for training and evaluation of machine learning models. 2023.

[KB15] D. P. Kingma and J. Ba. “Adam: a method for stochastic optimization.” International Conference on Learning Representations (2015).

[Kir+23] A. Kirillov, E. Mintun, N. Ravi, H. Mao, C. Rolland, L. Gustafson, T. Xiao, S. Whitehead,A. C. Berg, W.-Y. Lo, et al. “Segment anything”. IEEE/CVF International Conference on Computer Vision (2023).

[Lam+22] L. Lamm, R. D. Righetto, W. Wietrzynski, M. Pöge, A. Martinez-Sanchez, T. Peng, andB. D. Engel. “MemBrain: A deep learning-aided pipeline for detection of membrane proteins in Cryo-electron tomograms”. In: Computer Methods and Programs in Biomedicine (2022).

[Lam+24] L. Lamm, S. Zufferey, R. D. Righetto, W. Wietrzynski, K. A. Yamauchi, A. Burt, Y. Liu,H. Zhang, A. Martinez-Sanchez, S. Ziegler, et al. “MemBrain v2: an end-to-end tool for the analysis of membranes in cryo-electron tomography”. In: biorxiv (2024).

[LB21] Z.-L. Li and M. Buck. “Beyond history and “on a roll”: The list of the most well-studied human protein structures and overall trends in the protein data bank”. In: Protein Science 30.4 (2021).

[LH24] K. Lin and R. Heckel. “Robustness of deep learning for accelerated MRI: benefits of diverse training data”. International Conference on Machine Learning (2024).

[Liu+24] G. Liu, T. Niu, M. Qiu, Y. Zhu, F. Sun, and G. Yang. “DeepETPicker: Fast and accurate 3D particle picking for cryo-electron tomography using weakly supervised deep learning”. In: Nature Communications 15.1 (2024).

[LE22] T. Lüddecke and A. Ecker. “Image segmentation using text and image prompts”. IEEE/CVF Conference on Computer Vision and Pattern Recognition (2022).

[MS25] A. Martinez-Sanchez. “Template matching and machine learning for cryo-electron tomography”. In: Current Opinion in Structural Biology (2025).

[MSK24] V. J. Maurer, M. Siggel, and J. Kosinski. “PyTME (Python Template Matching Engine): A fast, flexible, and multi-purpose template matching library for cryogenic electron microscopy data”. In: SoftwareX 25 (2024).

[Moe+21] E. Moebel, A. Martinez-Sanchez, L. Lamm, R. D. Righetto, W. Wietrzynski, S. Albert, D. Lariviere, E. Fourmentin, S. Pfeffer, J. Ortiz, et al. “Deep learning improves macromolecule identification in 3D cellular cryo-electron tomograms”. In: Nature Methods 18.11 (2021).

[Oqu+23] M. Oquab et al. “DINOv2: learning robust visual features without supervision”. arXiv:2304.07193(2023).

[Pec+24] A. Peck, Y. Yu, J. Schwartz, A. Cheng, U. H. Ermel, S. Kandel, D. Kimanius, E. Montabana,D. Serwas, H. Siems, et al. “Annotating cryoET volumes: a machine learning challenge”. In:bioRxiv (2024).

[Per+18] E. Perez, F. Strub, H. d. Vries, V. Dumoulin, and A. Courville. “FiLM: visual reasoning with a general conditioning layer”. In: AAAI Conference on Artificial Intelligence 32.11 (2018).

[Rad+21] A. Radford, J. W. Kim, C. Hallacy, A. Ramesh, G. Goh, S. Agarwal, G. Sastry, A. Askell,P. Mishkin, J. Clark, et al. “Learning transferable visual models from natural language supervision”. International Conference on Machine Learning (2021).

[Ric+22] G. Rice, T. Wagner, M. Stabrin, and S. Raunser. TomoTwin demonstration dataset (1.0).2022. url: 10.5281/zenodo.7186070.

[Ric+23] G. Rice, T. Wagner, M. Stabrin, O. Sitsel, D. Prumbaum, and S. Raunser. “TomoTwin: generalized 3D localization of macromolecules in cryo-electron tomograms with structural data mining”. In: Nature Methods 20.66 (2023).

[RFB15] O. Ronneberger, P. Fischer, and T. Brox. “U-Net: convolutional networks for biomedical image segmentation”. Medical Image Computing and Computer-Assisted Intervention (2015).

[Sud+17] C. H. Sudre, W. Li, T. Vercauteren, S. Ourselin, and M Jorge C. “Generalised dice overlap as a deep learning loss function for highly unbalanced segmentations”. Deep Learning in Medical Image Analysis and Multimodal Learning for Clinical Decision Support (2017).

[TB20] M. Turk and W. Baumeister. “The promise and the challenges of cryo-electron tomography”.In: FEBS Letters 594.20 (2020).

[UVL16] D. Ulyanov, A. Vedaldi, and V. Lempitsky. “Instance normalization: the missing ingredient for fast stylization”. arXiv:1607.08022 (2016).

[Wag+19] T. Wagner, F. Merino, M. Stabrin, T. Moriya, C. Antoni, A. Apelbaum, P. Hagel, O. Sitsel, T. Raisch, D. Prumbaum, et al. “SPHIRE-crYOLO is a fast and accurate fully automated particle picker for cryo-EM”. In: Communications Biology 2.1 (2019).

[Wan+16] F. Wang, H. Gong, G. Liu, M. Li, C. Yan, T. Xia, X Li, and J. Zeng. “DeepPicker: a deep learning approach for fully automated particle picking in cryo-EM”. In: Journal of Structural Biology 195.3 (2016).

[Zen+23] X. Zeng, A. Kahng, L. Xue, J. Mahamid, Y.-W. Chang, and M. Xu. “High-throughput cryo-ET structural pattern mining by unsupervised deep iterative subtomogram clustering”. In: Proceedings of the National Academy of Sciences 120.15 (2023).

[Zha+24] Y. Zhao, H. Bian, M. Mu, M. R. Uddin, Z. Li, X. Li, T. Wang, and M. Xu. “Cryosam: training-free cryoet tomogram segmentation with foundation models”. International Conference on Medical Image Computing and Computer-Assisted Intervention (2024).

